# Learning with naturalistic odor representations in a dynamic model of the *Drosophila* olfactory system

**DOI:** 10.1101/783191

**Authors:** Ann Kennedy

**Affiliations:** Division of Biology and Biological Engineering, MC 156-29, Tianqiao and Chrissy Chen Institute for Neuroscience, California Institute of Technology, Pasadena, California 91125, USA

## Abstract

Many odor receptors in the insect olfactory system are broadly tuned, yet insects can form associative memories that are odor-specific. The key site of associative olfactory learning in insects, the mushroom body, contains a population of Kenyon Cells (KCs) that form sparse representations of odor identity and enable associative learning of odors by mushroom body output neurons (MBONs). This architecture is well suited to odor-specific associative learning if KC responses to odors are uncorrelated with each other, however it is unclear whether this hold for actual KC representations of natural odors. We introduce a dynamic model of the *Drosophila* olfactory system that predicts the responses of KCs to a panel of 110 natural and monomolecular odors, and examine the generalization properties of associative learning in model MBONs. While model KC representations of odors are often quite correlated, we identify mechanisms by which odor-specific associative learning is still possible.

## Introduction

Animals use their nervous system to translate signals from their sensory environment into appropriate behavioral responses. In some cases, these responses are hard-wired through genetic sculpting of neural circuits, such that certain stimuli drive behavioral responses in the absence of prior experience. But more often, responses to stimuli are modified over the course of an organism’s lifetime via associative learning, in which past experience is used to adaptively modify the neural circuits controlling behavior. An important feature of associative learning is that it should be stimulus-specific, meaning that a learned response to a reinforcer-paired stimulus will minimally affect the animal’s response to other unpaired stimuli. In cerebellum-like structures, this specificity is achieved via construction of a sparse, high-dimensional representation of stimulus identity, enabling learning of arbitrary associations by a read-out neuron via a simple synaptic learning rule^4-6^.

The mushroom body is a cerebellum-like structure that provides a neural substrate for odor-specific associative learning in insects^7-13^. The intrinsic cells of the mushroom body, Kenyon cells (KCs), form a sparse representation of odor identity^3,14,15^, while a small population of mushroom body output neurons (MBONs) receive convergent input from KCs, and show experience-dependent plasticity in their odor-evoked responses that underlies changes in behavior^8,17,18^. For learned associations in MBONs to be odor-specific, KC representations of odors must be not only sparse, but also high-dimensional, with low inter-odor correlations. Yet many odors evoke dense patterns of activation and high firing rates in olfactory receptor neurons (ORNs, the sensory neurons of the insect olfactory system), and inter-odor correlations in the ORN population are much higher than expected by chance^1,19^. Existing models of odor representations also neglect the dynamics of odor-evoked activity in the antennal lobe, which may substantially affect KC responses^20,21^. It is unclear how far these realities push mushroom body representations from the theoretical ideal.

To address this question, we present an olfactory system model capturing the connectivity and response dynamics of ORNs, projection neurons (PNs) and Kenyon cells (KCs), as well as the population activity of GABAergic local neurons (LNs) in the antennal lobe, and the GABAergic anterior paired lateral (APL) interneuron of the mushroom body (**Fig 1A**). This model uses observed connection probabilities between PNs and KCs, captures the temporal dynamics of PN input to the mushroom body, and models APL-mediated inhibition of KCs in a manner consistent with experimental data, together correcting inconsistencies of previous KC models with observed KC activity^20^. Model parameters at each stage of processing were fit to available experimental data, producing odor representations among model KCs that are consistent with the observed statistics of KC population activity.

**Figure 1.**
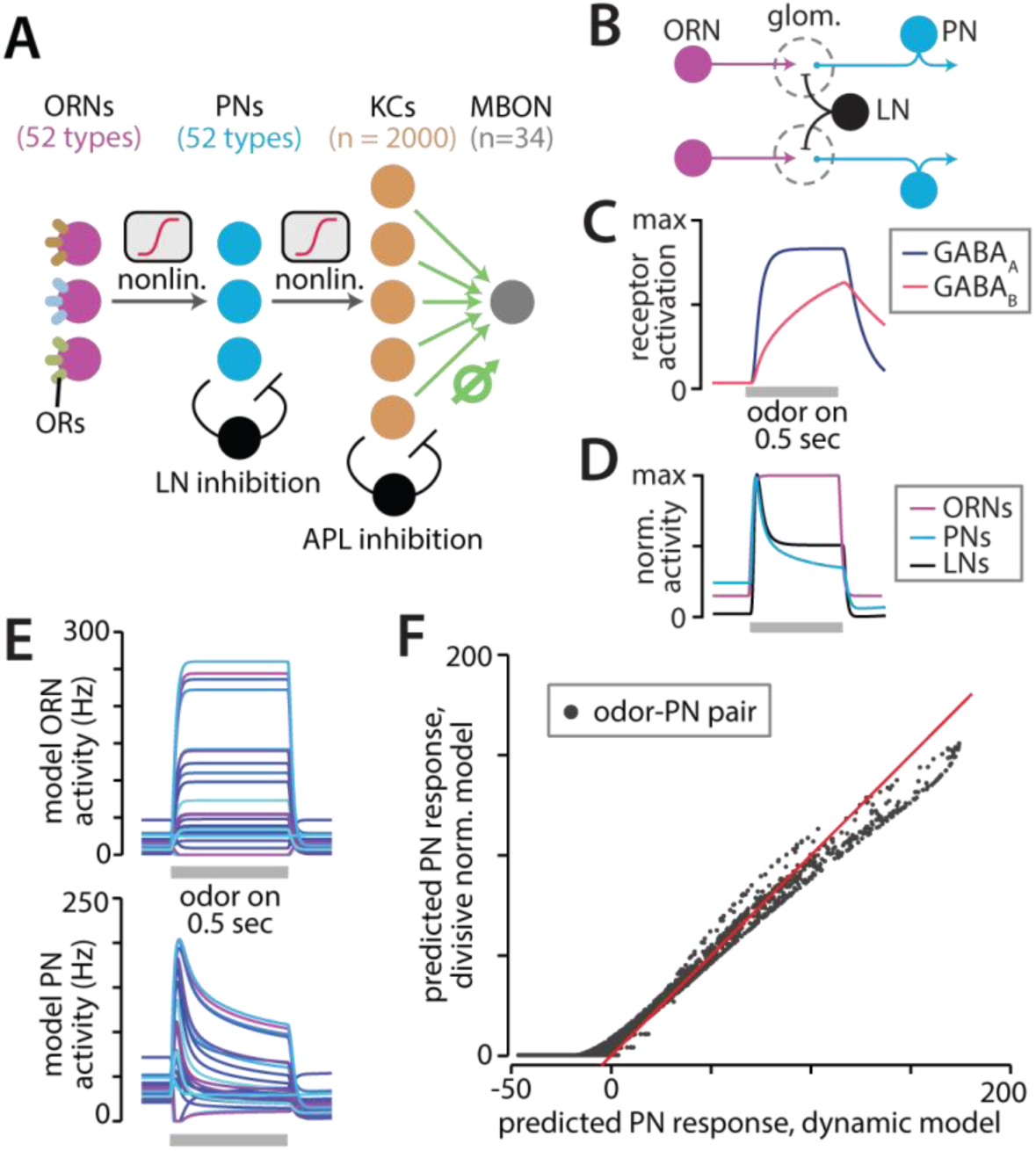
Building the dynamic PN model. **A)** Layers of the circuit model. Synapses from ORNs to PNs and from PNs to KCs are assumed to be fixed, whereas KC-MBON synapses (green) can be modified by learning. **B)** LNs innervate multiple glomeruli and have the net effect of normalizing odor representations across the PN population. **C)** Model dynamics of GABAA and GABAB receptor activation during a 0.5-second odor presentation. **D)** Population average activity in model ORNs, PNs, and LNs. LNs express GABAA receptors only, while PNs express both GABAA and GABAB receptors, leading to the difference in response shape between the two populations. **E)** Example responses of 23 model ORNs and their cognate PNs to one example odor (isopentyl acetate) from the 110-odor dataset. **F)** Comparison of predicted time-averaged PN responses in the static model^2^, and in the model proposed here. Each point is the response of one of 23 PNs to one of 110 odors.

Strikingly, our model predicts a much higher degree of overlap in KC representations than is expected in a model that assumes independent and uncorrelated odor representations by ORNs. As a result, our model predicts that associations learned by a model MBON will lead to changes in the MBON’s response to other, untrained odors, an effect akin to overgeneralization. However, we identify two mechanisms by which odor overlap or overgeneralization may be reduced. First, the addition of developmental homeostatic plasticity in the KC population substantially reduces odor overlap in model KCs, thus reducing the rate of overgeneralization during learning. And second, a modification of the associative learning rule can allow the model MBON to produce numerous odor-specific learned responses even without the need for homeostatic plasticity in KCs. Each mechanism makes testable predictions regarding odor representations in KCs and the dynamics of MBON responses during learning. By linking together experimental data from each stage of insect olfactory processing, our model thus identifies potential mechanisms for improved odor learning, and serves as a tool for more focused design of future experiments investigating learned odor-guided behaviors.

## Results

### A firing rate model of PN activity closely fits prior models of PN response normalization

Input to our model is generated at the level of ORNs, and was sampled from experimentally observed responses of 23 odorant receptor (OR)-expressing ORNs to 0.5-second presentations of each of a panel of 110 odors^1^. Odor representations are then relayed from ORNs to PNs to KCs, thus undergoing two successive steps of nonlinear transformation. The first of these occurs in the antennal lobe, where a population of GABAergic local neurons (LNs) receives input from ORNs and PNs, and provides feedback inhibition onto ORN terminals^22,23^ (**Fig 1B**). While multiple subtypes of LNs have been described, the LN population shows largely the same spatial pattern of activation regardless of odor^24^, and the primary effect of this inhibition is to divisively normalize the PN population response. Normalization makes odor representations in PNs more invariant to odor concentration, and reduces the dynamic range of PN input to the mushroom body^2,24-27^.

Previous models of PN response normalization were fit to the time-averaged firing rate of PNs in response to odors. But experimental characterization of inhibition in the antennal lobe has shown that it is slow compared to excitation^21^; this inhibition, combined with synaptic depression, contributes to a transient peak in odor-evoked PN firing rates at odor onset that decays to a lower, steady-state value as inhibitory currents kick in^25^ (**Fig 1C**). To investigate the impact of this transient peak in PN activation on odor representations, we simulated PN response normalization in a dynamic model of the antennal lobe.

We determined the dynamics of LN inhibition by fitting time constants of GABA binding to GABAA and GABAB receptors, using previously published IPSP recordings in LNs and PNs^21^ (**Fig 1D**; see Methods). We fit model parameters—an ORN-LN nonlinearity, and two constants determining the strength of inhibition—to minimize the difference between time-averaged PN responses in our dynamical model, and PN responses predicted by a previously published (but non-dynamical) model of the ORN-PN nonlinearity, which we call the “static” model^2^. The time-averaged responses of our fit dynamical model were closely matched to PN responses in the divisive normalization model for all odor/PN pairs (**Fig. 1F**).

### Slow lateral inhibition shapes response dynamics in PNs and normalizes sustained, but not initial, glomerulus activity

We next contrasted the dynamics of PN responses in our model with the dynamics of experimentally recorded PNs. Model PN responses to a “narrow” odor (one that only strongly activates a single glomerulus) become more transient when that odor is mixed with a weak or strong “broad” odor (one that activates many glomeruli), an effect that has been reported previously (**Fig 2A**)^2^. This result was obtained without any additional adjusting of model parameters. We also simulated the response of model PNs to experimentally measured ORN responses evoked by a panel of 10 chemical odors, each presented at dilutions of 1e-8, 1e-6, 1e-4, and 1e-2, as well as by a panel of 8 fruit extracts, presented at dilutions of 1e-6, 1e-4, 1e-2, and undiluted^1^. We found that our dynamical model reproduced another experimental observation of PNs, namely that PN responses become more transient at higher odor concentrations^25^ (**Fig 2B**). The time-averaged PN population firing rate increases only marginally across a four order of magnitude change in odor concentration, consistent with the previously reported concentration invariance of odor representations in PNs compared to ORNs (**Fig 2C**; **Fig 2D** dark blue line), as is seen in vivo^2,20,27^. However, peak PN firing rates in our model show a concentration dependence close to that of ORNs (**Fig 2D**, light blue line), due to the slow kinetics of GABA receptor activation in model PNs. This transient peak in PN firing rates has consequences for the concentration dependence of KC responses to odors, as we will later discuss.

**Figure 2.**
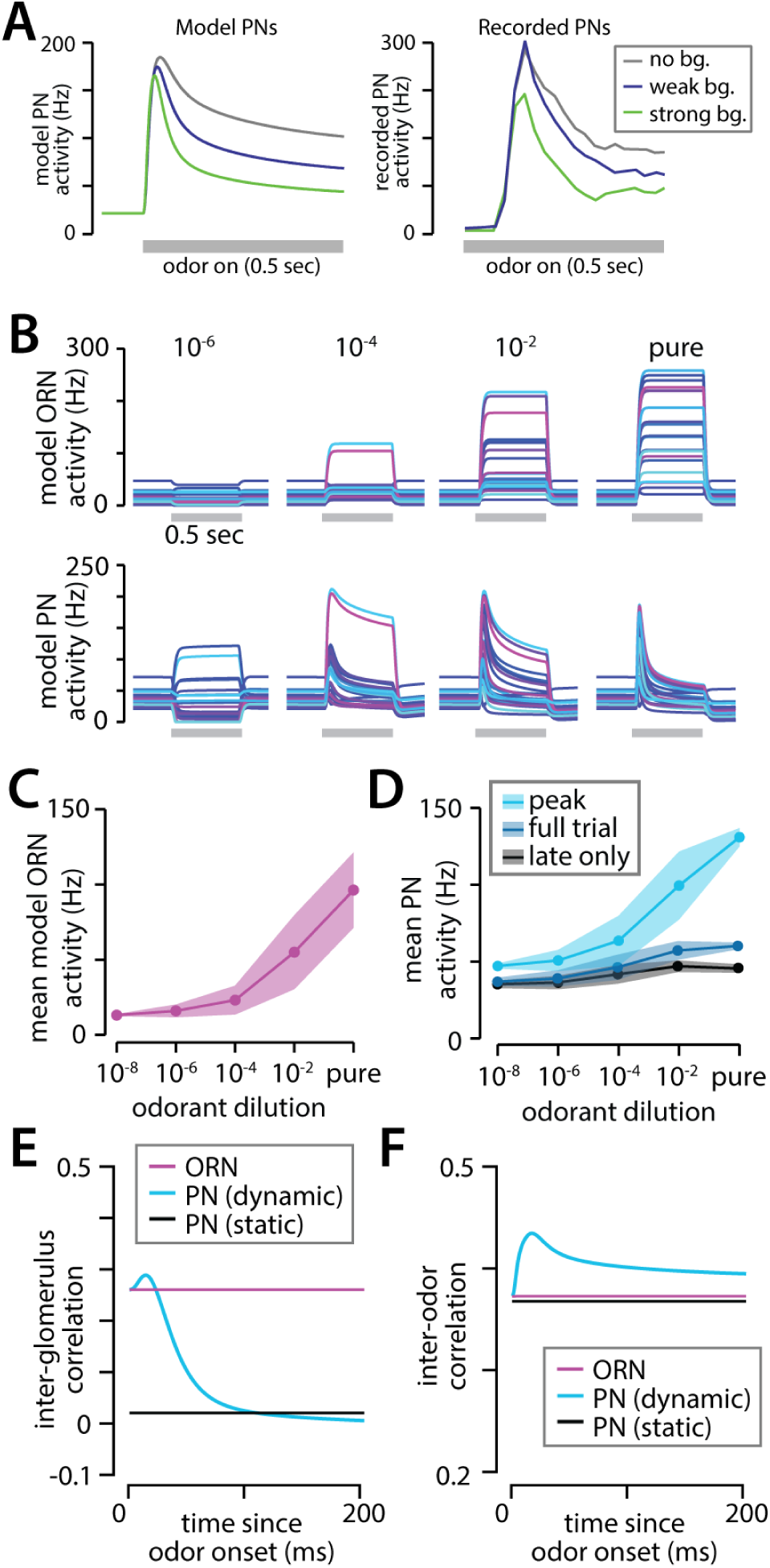
PN representations in the dynamic model. Response of a PN to a narrow odor that strongly activates its cognate ORN. Mixing the narrow odor with a broad odor causes PN response to become more transient as broad odor concentration is increased, due to lateral inhibition from other glomeruli. **Left:** simulated PN activity, **Right:** experimental data, reproduced from ^2^. **B)** Responses of model ORNs (top row) and PNs (bottom row) to 0.5-second presentations of multiple concentrations of banana extract. At higher concentrations, PN responses become more transient due to increased LN recruitment. **C)** Population-averaged firing rate of model ORNs in response to ten chemical odors at dilutions 10^-8^, 10^-6^, 10^-4^, 10^-2^, and eight fruit odors at dilutions 10^-6^, 10^-4^, 10^-2^, and undiluted. ORN population activity increases at higher odor concentrations. **D)** Population averaged firing rate of model PNs. The time-averaged PN response shows little concentration dependence (“full trial”), as reported previously. In contrast, activity during the initial peak of the PN response much more resembles the concentration dependence of ORNs (“peak”) This high initial concentration dependence is countered by a very low concentration dependence by the end of the stimulus interval (“late only”). **E)** The average correlation between pairs of glomeruli as a function of time, across the 18 odors tested in A-C, 10^-2^ dilution. Odor representations taken to be the instantaneous firing rate of model neurons, in Hz. Divisive normalization decorrelates PN representations in the static model, but does not affect the initial transient response of PNs in the dynamic model. **F)** Average correlation between glomerular representations of pairs of odors (same odors as in D), shows no difference between ORNs and PNs in the static model, and a slight increase in the dynamic model.

### Lateral inhibition decorrelates glomeruli, but not odors

Previous work has shown that divisive lateral inhibition between glomeruli serves to decorrelate the activity of PNs over a set of presented odors^2^; we will call this the inter-glomerular (or inter-ORN or –PN) correlation. In the static model, inter-PN correlations are substantially lower than inter-ORN correlations, as reported previously (**Fig 2E**, black line). But in the dynamic model, the slow kinetics of lateral inhibition cause inter-PN correlations to change over time: PN representations are as correlated as their cognate ORNs at odor onset, when PN firing rates are also highest; it is only at 100ms after odor onset that inter-PN correlations in the dynamic model approach those of the static model (**Fig 2E**, blue line).

In addition to measuring correlations between pairs of glomeruli, we also consider the effect of lateral inhibition on correlations in the PN population responses between pairs of odors (inter-odor correlations). Strikingly, there was little difference between inter-odor correlations at the level of ORNs and PNs in the static model, while the dynamic model even showed a slight increase in inter-odor correlations (**Fig 2F, Fig S1**). This finding demonstrates the important fact that inhibitory normalization can produce very low inter-glomerular correlations, without having an effect on inter-odor correlations. Yet inter-odor correlations are an important statistic to consider for the purpose of olfactory computation: excluding differences in total response magnitude, inter-odor correlations are a good predictor of the discriminability of an odor pair, while inter-glomerular correlations are not. We therefore do not expect LN normalization of PNs to have a strong impact on odor discriminability at the level of KCs.

### Incorporating PN input into a spiking model of the mushroom body

We next used the dynamic PN model as input to a model of mushroom body itself, to investigate the transformation of odor representations from PNs to KCs. The mushroom body is composed of approximately 2000 KCs, each of which has a small number of dendritic claws (6.8+/-1.7, mean ± STD, N=200 KCs; data from ^16^). Each KC claw forms a synapse with a single PN in a random and independent manner, meaning that the probability of PN-KC synapse formation is independent of KC location, type, and of other inputs to the same KC^16,20^.

We modeled KCs as a population of 2000 leaky integrate-and-fire neurons. Each model KC was assigned a number of dendritic claws, randomly sampled from a set of 200 observed KC claw counts^16^. For each claw, we then randomly selected an input glomerulus, with probability of selecting each glomerulus taken from a set of 342 observed PN-KC synapses in the mushroom body of adult *Drosophila*^16^ (**Fig 3A**). Interestingly, these data show that some PNs have more axonal projections to the mushroom body than others, and are more likely to form synapses with KCs. We found that the observed PN-KC connection probabilities are correlated with spontaneous firing rates of each PN’s cognate ORN^1^, suggesting that spontaneous PN activity might shape the formation or retention of PN-KC synapses (**Fig 3B, D**). In contrast, mean odor-evoked PN firing rates were not correlated with PN-KC connectivity (**Fig 3C, E**).

**Figure 3.**
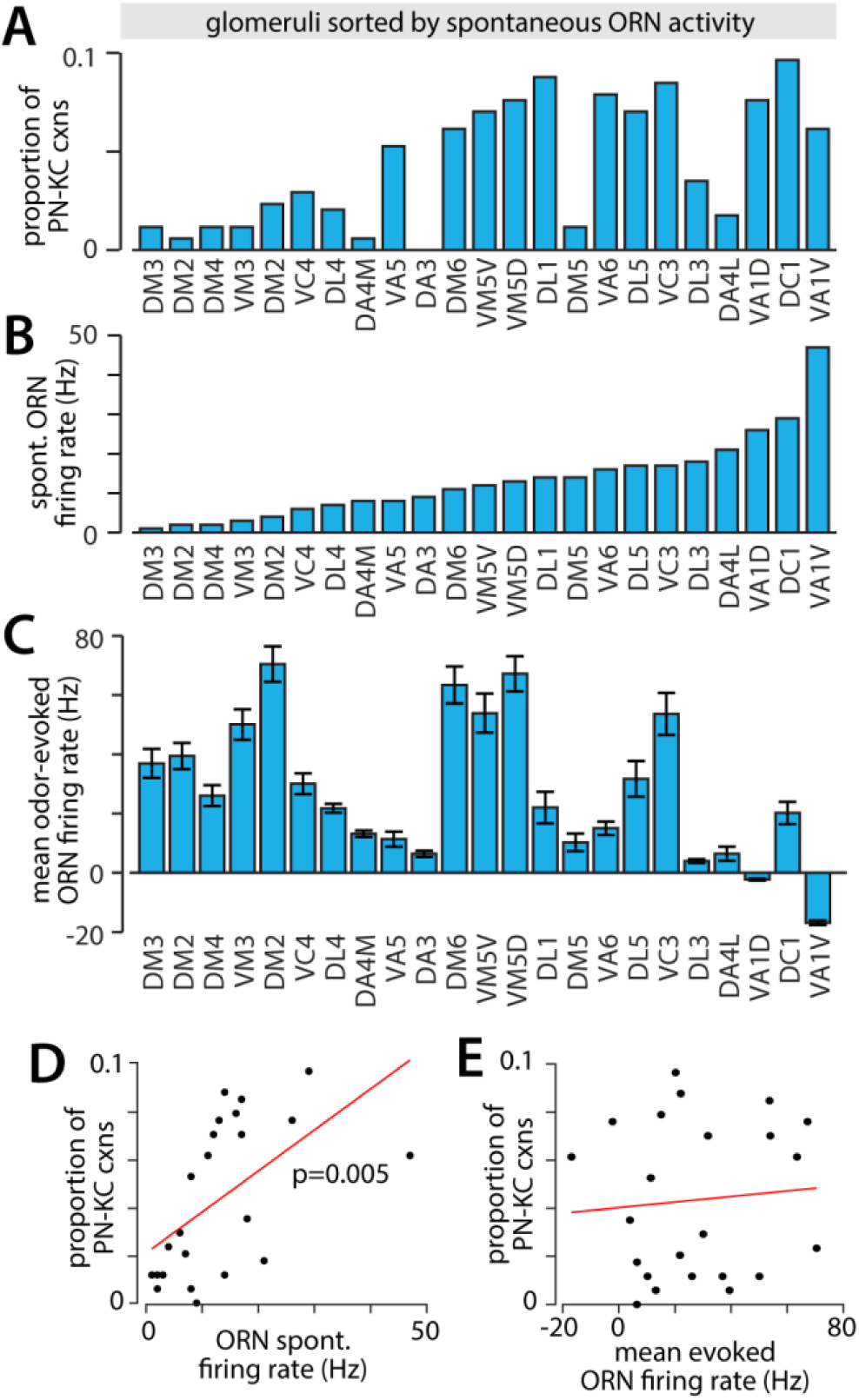
Non-uniform properties of PNs. Glomeruli in A-C are sorted by the spontaneous firing rate of ORNs as reported by ^1^. **A)** The fraction of PN-KC synapses derived from each of 23 glomeruli, as measured in ^16^ (PNs not included in the Hallem and Carlson dataset have been excluded.) **B)** Spontaneous ORN firing rates from ^1^. **C)** Odor-evoked ORN firing rates from^1^, for 110 tested odors, mean±SEM. D) Statistically significant correlation between the spontaneous firing rates of ORNs, and the probability of those ORNs’ cognate PNs forming synapses with KCs. Slope of a linear fit (red) is significantly different from value expected by chance (p=0.00546). Each point is one glomerulus, same data as in **A-B. E**) No correlation is observed between mean odor-evoked firing rates of ORNs and probability of PN-KC synapse formation. Data from panels **A** and **C**.

In addition to feedforward input from PNs, KCs receive input from the anterior paired lateral neuron (APL), a single massive GABAergic interneuron^28^. APL innervates the entirety of the mushroom body, and exhibits graded release of GABA at a level proportional to the amount of KC activation^29^. In a previously published model of the mushroom body^20^, APL was assumed to decorrelate KC representations by removing the first principal component of PN responses to the 110-odor ensemble; while this approach substantially improves mushroom body performance on odor learning tasks, it results in patterns of KC activity that do not match well with available data, for example predicting more KCs would respond to narrow odors that only activate a single glomerulus than to broad odors that activate many glomeruli. In the absence of detailed data on APL-KC connectivity, we assumed that APL receives equal input from all KCs, and divisively inhibits all KCs with equal strength. Because an average of 10% of KCs respond to a given odor, and silencing of APL has been found to roughly double this number^30^, we set KC thresholds to achieve 20% odor sparsity, then uniformly scaled synaptic weights from APL to KCs and from KCs to APL to reduce sparsity to 10% (see Methods). Finally, all KCs were assumed to have the same activation threshold relative to their average amount of spontaneous PN input, though we revisit this assumption in the next section.

We first examined the lifetime sparsity of model KCs across a panel of 25 test odors, selected for their chemical similarity to an odor panel from a previous study of KC sparsity^3^, and computed lifetime sparseness of KCs and of odors as in ^13^. Lifetime sparseness varies between 0 and 1 (1=sparsest). The distribution of lifetime KC sparseness in the model closely matched that of experimental data (**Fig 4A**), and demonstrates the increased sparseness of odor representations in the mushroom body compared to the activity of projection neurons (**Fig 4B**). We similarly computed the lifetime sparseness of tested odors among both KCs and PNs, and found that most odors in the model KC population activate a small proportion of KCs, as was seen in the data (**Fig 4C-D**).

**Figure 4.**
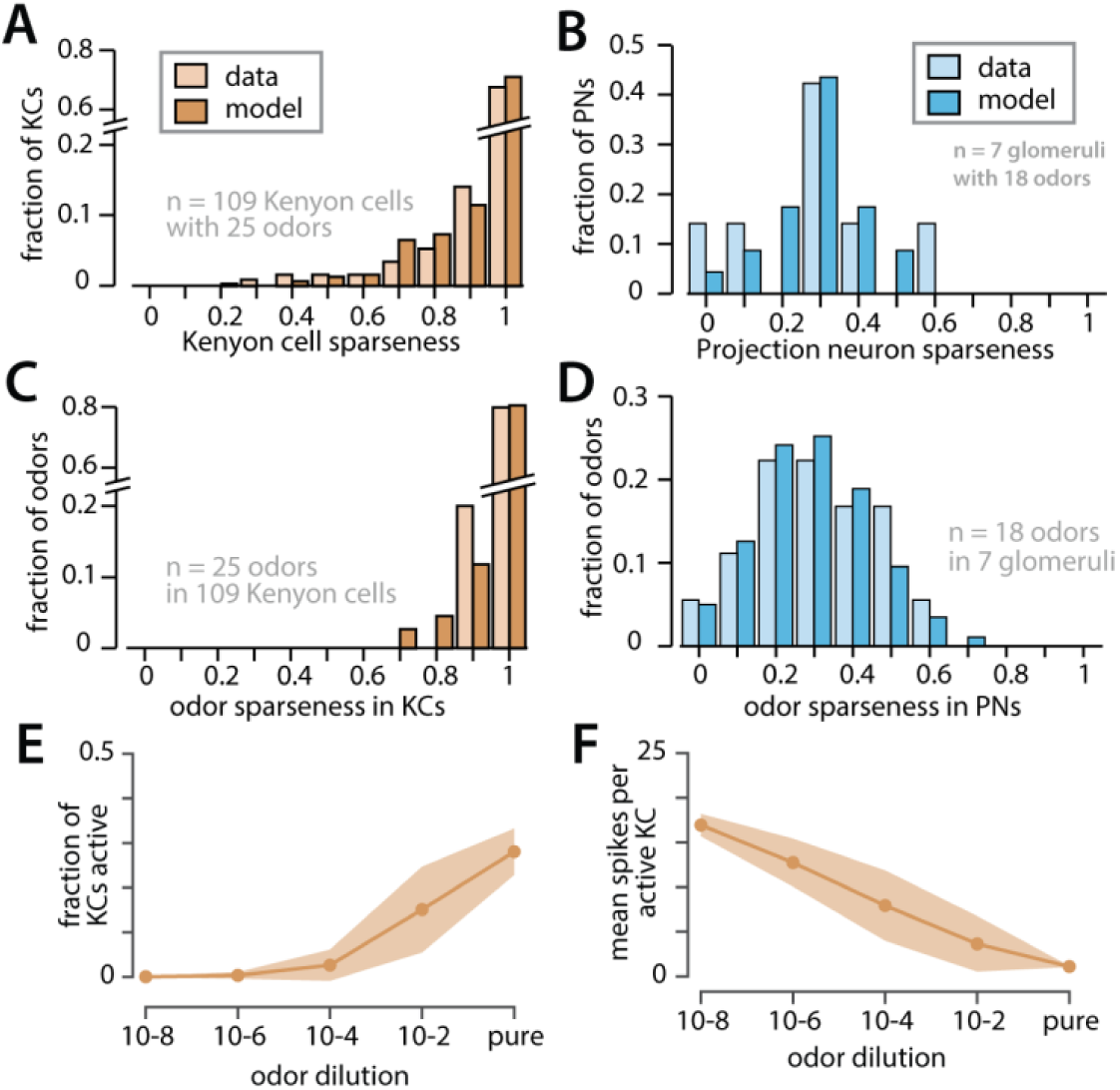
Sparseness properties of model KC population. **A)** Lifetime sparseness of 109 model (orange) or experimentally observed (tan) Kenyon cells, using a panel of 25 test odors. Experimental data from ^3^. **B)** Lifetime sparseness of 7 model or experimentally observed PNs, showing far denser responsivity to odors. Experimental data from ^3^ **C)** Odor representation sparseness of a panel of 25 test odors in a population of 109 KCs (same data/simulation as in **A). D)** Odor representation sparseness among PNs. E) Fraction of model KCs activated by different odor dilutions, using PN input from same dataset as **Figure 2C-D. F)** Average number of odor-evoked spikes in KCs that responded to different odor dilutions, showing a decrease in KC response persistence at higher odor concentrations.

Using the odor dilution dataset, we then investigated odor dependent activity of our model KC population. The number of model KCs responding to an odor increases with odor concentration (**Fig 4E**), however interestingly the average number of spikes per KC decreased (**Fig 4F**). Thus, as in PNs, an increase in odor concentration caused model KC activity to become more transient, while the number of KCs responding to an odor increased. Because of this, the time-averaged PN response to odors was a poor predictor of the number of KCs responding to an odor, while the peak height of the PN population response was a much better predictor (**Fig 4E-F**, compare to **Fig 2C-D**). Similarly, and in contrast to a previous model^20^, our model predicts more active KCs, but fewer spikes per KC, for broad odors compared to narrow odors (**Fig S2**).

While it has previously been shown that PN responses become more transient at higher odor concentrations^2^, it is unknown whether this effect is seen in KCs. If experimental observations show that KCs do not show sustained firing in the presence of odors, our mushroom body model could be updated to include synaptic depression at PN-KC synapses, making KC responses to weaker odors more transient and reducing the concentration dependence seen in **Fig 4F**.

### Homeostatic plasticity reduces overlap of odor representations by KCs

To estimate the degree of overlap of odor representations in the model KC population, we simulated KC responses to the 110-odor dataset and computed the Intersection Fraction (IF) for all odor pairs, defined as the percent of KCs responding to odor *a* that also responded to odor *b* (this metric is a good predictor of generalization during learning, see following sections and **Fig S5**). To distinguish between contributions of KC threshold and recurrent inhibition to odor representations, we simulated KC responses in models with and without APL inhibition; in models without APL inhibition, KC thresholds were elevated to preserve an average of 10% of KCs responding per odor. Across all tested versions of the model, APL inhibition decreased the number of “silent” odors (odors activating no KCs) compared to the no-APL model in which sparseness was achieved by elevating KC thresholds (**Fig 5C**).

**Figure 5.**
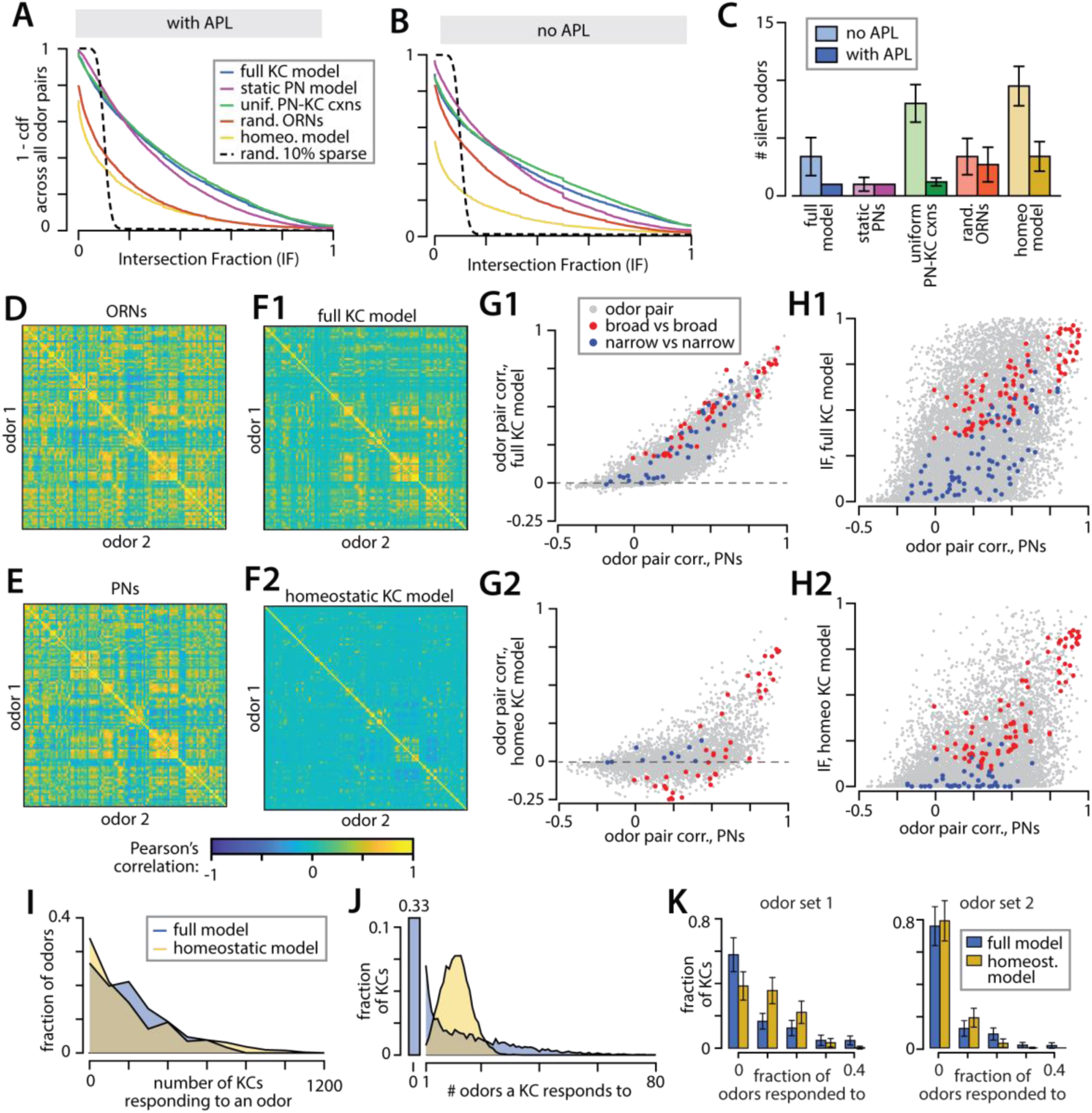
Degree of overlap in odor representations for several versions of the mushroom body model. **A)** Distribution of Intersection Fraction (IF) for all odor pairs, in multiple versions of the mushroom body model and for random 10% sparse vectors. **B)** Plots as in A, but for models incorporating inhibitory normalization by APL. C) Average number of odors that failed to activate any KCs for each model, with and without inhibition (colors as in **A). D**) Pearson’s correlation between population responses to pairs of odors in model ORNs **E)**, Same in PNs **F1-2)** Same in KCs from the full (**F1**) and homeostatically tuned (**F2)** models. **G1-2**) Scatter plot of Pearson’s correlation data from PNs vs KCs in the full (**G1**) or homeostatically tuned **(G2**) models. Correlation between the 10 broadest odors is shown in red; between the 10 narrowest odors in blue. Same data as in **E-F**; legend as in H. **H)** Pearson’s correlation in PNs vs IF in KCs in the full (**H1**) or homeostatically tuned (H2) models. **I)** Distribution of the number of model KCs responding to each odor in the Hallem and Carlson dataset, in both the full KC model and the homeostatically tuned model. **J)** Distribution of the number of odors each KC responded to, for the full and homeostatically tuned KC models. Bar to the left is the fraction of cells in the original model that did not respond to any of the 110 odors. **K)** Fraction of odors model cells responded to, for two sets of mostly broad (left) and mostly weak/narrow (right) odors. All panels computed from ten repeated simulations (generation of PN-KC connectivity matrices); inter-simulation variability was low (see **Fig S3**.)

If each odor were to activate a randomly selected 10% of KCs, the mean IF across odor pairs would also be 10% (**Fig 5A-B**, black dashed line). While this is close to the mean IF of our full mushroom body model, the distribution of IFs observed in our model is much broader: 58% of odor pairs in the model with APL inhibition had an IF of 20% or more, compared to less than 0.01% expected given random 10%-sparse representations. We find that representation overlap arises primarily from correlations in odor representations among ORNs: shuffling ORN responses across odors and glomeruli decreased the proportion odor pairs with IF>20% from 58% to 27% (**Fig 5A-B**, red line). In contrast, giving all PNs equal probability to form synapses with KCs had little effect on overlap (**Fig 5A-B** green line), nor did replacing the dynamic PN model with the original static PN model (**Fig 5A-B**, purple line).

Without changing the PN inputs to the mushroom body, the overlap between odor representations might be reduced by tuning individual KC thresholds or by modifying the inhibitory APL feedback. We tested a “homeostatic” rule to set the thresholds of individual KCs, both in the presence and absence of APL inhibition. For each model KC, we computed its PN input for a subset of odors in the Hallem and Carlson dataset, then set that KC’s threshold so that it responded to 10% of tested odors. This rule substantially decreased the Pearson’s correlation and IF between KC representations of odors (**Fig 5D-H**), and also substantially decreased the overlap in odor representations (**Fig 5A-B**, yellow line). Interestingly, the IF of odor representations among KCs was only modestly correlated with the correlation between odor pairs among PNs (**Fig 5H**). However, we did find that pairs of narrow odors typically had a low IF, while broad odors were more likely to have a high IF.

While the proportion of KCs activated by a given odor is similar in the original and homeostatically tuned models (**Fig 5I**), the original model produces a large number of “silent” KCs (33%) that did not respond to any of the 110 odors, as well as many KCs that respond to many odors (**Fig 5J**). This suggests that it should be possible to determine which model is a better fit to the mushroom body by computing the distribution of KC response sparsities across a panel of presented odors. In designing such an experiment, we found that a test panel of odors are best suited to distinguishing models. Although the two models showed different response distributions to random subsets of 25 broad odors (selected from a set of aldehydes, ketones, aromatics, and alcohols in the 110-odor dataset) (**Fig 5K**, odor set 1), a second odor panel of amines, lactones, and acids showed little difference between models (**Fig 5K**, odor set 2).

We also observe a modestly negative Pearson’s correlation between the KC population representations of alcohols vs some esters in the homeostatic model, but not in the original model (**Fig 5F-G, Fig S3**). A search for similarly anticorrelated odor representations in the mushroom body might provide further support for the homeostatic model of KC responses.

### Rapid overgeneralization of odor-evoked responses by a naïve learning model

Having built a model that takes into account the dynamics and statistics of PN input to the mushroom body, we next investigated the suitability of our model KCs as a basis for odor-specific associative learning by MBONs. Plasticity at KC-MBON synapses is modulated by a population of dopaminergic neurons^31^, different subsets of which are activated by multiple rewarding^32,33^ or punishing^34^ stimuli, or by the animal’s behavioral state^35^. We expect associative learning to be odor-specific, meaning that learning modifies the response of an MBON to a set of reinforcer-paired odors, while minimally altering that MBON’s response to other “unpaired” odors. This reflects our expectations of the specificity of learned behavior: learning to associate one or more odors with a reward or punishment should not drastically change an animal’s response to other, different odors (although see Discussion.) We also expect learning to be possible with incomplete prior sensory experience: an animal should be able to learn to associate one or more odors with a reward or punishment without also being given the identity of all other odors that are *not* associated with that reward or punishment. This can be seen as a one-class classification problem, in contrast to the classical discrimination problem of learning from both positive vs negative examples.

If the neural representation of stimuli is high-dimensional, a single-layer perceptron performs quite well in one-class classification^36^. We used the perceptron algorithm to train a model MBON to respond to a randomly selected subset of odors, reflecting the set of reinforcer-paired odors. We then tested the specificity of learned responses by computing the fraction of unpaired odors to which the MBON also responded after learning, which we call the MBON’s probability of overgeneralization (**Fig 6A**). As the number of paired odors increases, the probability of overgeneralization by the MBON increases at a rate determined by the degree of overlap of odor representations. If KC representations of odors are taken to be random, 10% sparse binary vectors, a large number of associations (∼20) can be learned by a single MBON before overgeneralization sets in (**Fig 6B**, black line). In contrast, the high correlation of model KC responses to actual odors leads to rapid overgeneralization of associative learning after a few odor pairings (**Fig 6B**, blue line). Overgeneralization was higher for training on broad odors than for training on narrow odors (**Fig S4**), and for single trained odors, the probability of overgeneralization to an unpaired odor was well predicted by the percent of KCs responding to that unpaired odor that also responded to the trained odor (**Fig S5**). Thus, overgeneralization can be seen as a function of the overlap in odor representations among KCs. Consistent with this finding, we could substantially reduce the rate of overgeneralization by decorrelating odor representations by shuffling ORN identities across odors, or by substituting representations from the homeostatic KC model (**Fig 6B, Fig S5**). However even in these cases, overgeneralization by the model MBON is much higher than in the idealized, 10%-sparse case.

**Figure 6.**
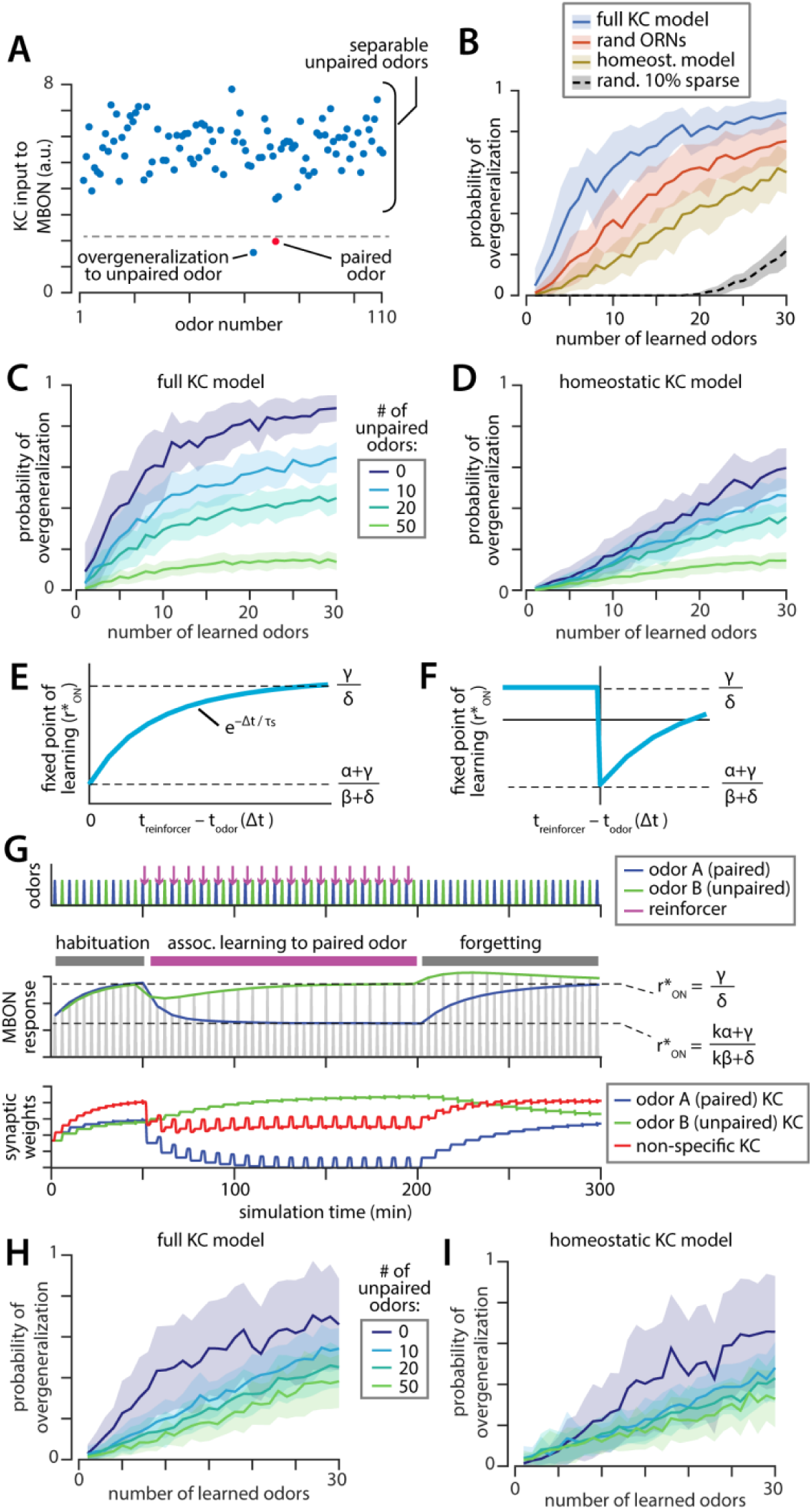
Learning odor-specific associations. **A)** Cartoon example MBON response to 110 odors, showing one paired odor whose representation has become depressed following pairing with a reinforcer (red) and a “overgeneralized” unpaired odor. **B)** Probability of MBON overgeneralization to unpaired odors after learning multiple odor associations by the same MBON using a single-class perceptron algorithm, for random 10% sparse vectors (black) and for several versions of the KC model. (Envelopes show mean±STD over 10 model replicates.) **C-D)** Probability of overgeneralization to novel unpaired odors decreases when the perceptron algorithm is allowed to learn from examples of both paired and unpaired odors. **E)** The fixed point of the two-part, eligibility-trace-based learning rule is a function of learning rule parameters *α, β, γ, δ*, the decay constant of the synaptic eligibility trace *τ*_*s*_, and the delay between odor and reinforcer presentation *Δt*. **F)** For long inter-trial intervals, the eligibility-trace-based learning rule can produce bidirectional changes in the MBON response to odors. **G)** Dynamics of MBON responses during pairing of an odor with dopamine. Top row: timing of odor and reinforcer presentations. Middle row: magnitude of MBON response to paired and unpaired odors. Bottom row: dynamics of KC-MBON synaptic weights, for KCs that responded specifically to the paired or unpaired odor, or for KCs that responded to both odors (red). Note compensatory change in synaptic weights from the odor B KCs during learning, in response to decreased synaptic weights from the nonspecific KCs. **H-I)** Probability of MBON overgeneralization to novel unpaired odors using the two-part learning rule, using the full and homeostatic forms of the KC model population.

### Learning from incomplete negative examples reduces overgeneralization to novel odors

Learning does not occur in a vacuum: in addition to odors that are predictive of reward or punishment, flies encounter many environmental odors that bear little to no predictive power. Learning to “ignore” a small number of unpaired odors may help the fly to avoid overgeneralization in the MBON response to paired odors. We tested this idea by training the perceptron algorithm on a two-class classification problem, with a subset of unpaired odors present during learning. Overgeneralization was then tested on the remaining unpaired odors. Inclusion of a relatively small number of unpaired odors substantially reduced the rate of MBON overgeneralization using the full KC model, while overgeneralization was more mildly reduced in the homeostatic model (**Fig 6C-D**). Learning to ignore unpaired odors therefore seems to achieve a similar effect to our proposed homeostatic plasticity among KCs, and may provide an alternative means of achieving odor-specific responses given correlated representations of odors by the KC population.

### Achieving odor-specific associative learning in the dynamic model

Motivated by our findings with the perceptron model, we designed a synaptic plasticity rule that would allow a model MBON to form odor-specific associative memories while also learning to ignore odors not associated with reward. We begin by considering an anti-Hebbian associative learning rule of the form 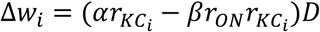, where *D* is nonzero in the presence of dopamine and *α* and *β* are nonnegative scalars. Repeated pairing of an odor with dopamine in this model leads to a reduction in KC input to the MBON from all KCs that responded to the paired odor. We chose an anti-Hebbian rule because synaptic weights under this rule converge to a stable fixed point at *r*_*ON*_ = *α*/*β*, however we also considered purely associative or depressive learning rules of the form 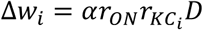, with bounded synaptic weights to impose stability.

To enable learning in the presence or absence of dopamine, we added a second term to our learning rule, giving it the form 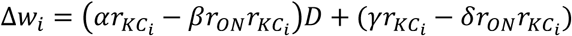. For a given value of *D*, this rule has a single stable fixed point *r*_*ON*_ = (*γ* + *Dα*)/(*δ* + *Dβ*). If *D* is taken to be 1 in the presence of reinforcement, and 0 otherwise, this rule therefore pushes *r*_*ON*_ to the stable fixed point *γ*/*δ* for unpaired odors, and to the stable fixed point (*γ* + *α*)/(*δ* + *β*) for paired odors.

Finally, to address the fact that odors and rewards are often not presented at the same time, we converted our rule to a time-dependent form by inclusion of a synaptic eligibility trace for plasticity^37^ 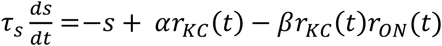, with which the KC-MBON synaptic weight is determined by learning rule 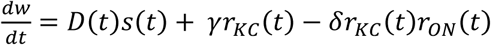. In this form, the fixed point of learning becomes 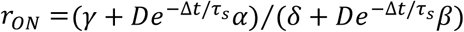, where Δ*t* is the mean time delay between odor and reinforcer presentation, and *τ*_*s*_ is the decay rate of the synaptic eligibility trace *s*. Because the fixed point is dependent on Δ*t*, this rule can produce both potentiation and depression depending on the timing of reinforcement and odor presentation (**Fig 6E-F**), although only one direction of change (here depression) is dopamine-dependent.

While many rules of this form exist, their most salient common feature is that when dopamine reinforcement is absent (D=0), KC-MBON synaptic weights will still evolve following odor presentation (**Fig 6G**). This is consistent with previous observations of MBON habituation to familiar odors^18^, and also with experimental characterization of a reinforcer-modified STDP rule identified in KC-MBON synapses of locust, where the shape of the rule is modulated by the presence of octopamine^38^.

To investigate overgeneralization by our two-part learning rule, we interspersed reinforcer-paired trials with unpaired trials using other odors, setting the dopamine signal D=1 on reinforcer-paired trials and D=0 otherwise (an alternative form of this model, in which the dopamine signal is more complex, is presented in the Methods). The two-part learning rule substantially reduced the rate of learning overgeneralization, even when the number of unpaired odors was small (**Fig 6H-I**). Because the fixed point of this learning rule is set by parameters *α, β, γ*, and *δ* that are characteristic of a given MBON, this model predicts that the MBON response after learning will be independent of learned odor identity, and will be the same for all odors paired with reinforcement. This model also predicts that inhibiting dopaminergic neurons after learning, or stopping reinforcement, will lead the MBON response of a previously reinforced odor to revert back to the unpaired odor fixed point.

## Discussion

We have presented a dynamic model of olfactory representations as they are relayed from the olfactory periphery to the mushroom body, and shown how such a circuit can underlie odor-specific associative learning of naturalistic odors by MBONs. While the architecture of the mushroom body can produce sparse and high-dimensional odor representations in the idealized case of independent and uncorrelated odors^6^, it is unknown to what extent this holds for naturalistic odor representations. We present a dynamic model of the insect olfactory system as odor representations are relayed from ORNs to PNs to KCs, and model its responses to a panel of 110 monomolecular odors^1^. Our model reproduces experimentally observed dynamics of PN responses, allowing us to estimate the extent of odor normalization in antennal lobe input to the mushroom body. We use our dynamic PN model as input to a population of spiking model KCs, whose activity is normalized by a revised model of the GABAergic neuron APL^29,30^. The model at all stages is constrained by available experimental data, although we note instances (such as details of APL-KC connectivity and the setting of KC spiking thresholds) where further information could be used to refine our model.

In a first instantiation of the KC model, we found that correlations between odor representations are still present in the model KC population, and that these correlations result in overgeneralization in a model of associative odor learning. Adding a homeostatic threshold-tuning model substantially decorrelated odor representations in the model KC population, and reduced the rate of overgeneralization. While homeostatic plasticity of KCs has not been reported to our knowledge, our observation that spontaneous ORN firing rates were correlated with the probability of their cognate PNs forming synapses with KCs does suggest that activity-dependent plasticity may contribute to the wiring of the mushroom body: further investigation of the effect of developmental environment on PN-KC connectivity could test this hypothesis. We also suggest imaging experiments targeting specific panels of odors, or experiments testing for anti-correlation of certain odor pairs, that would support the homeostatic model.

We also identified a class of learning rules by which a model MBON could learn multiple odor-reinforcer pairings without overgeneralizing to unpaired odors even without homeostatic plasticity among KCs. Central to our learning rules is the inclusion of a learning process that occurs in the absence of a paired reinforcer (dopamine), reducing the contribution of “non-specific” KCs (KCs that respond to both paired and unpaired odors) to the learned MBON response. Our two-part learning rule is consistent with evidence of MBON habituation to repeated odor presentation^18^, and suggests that this observed habituation may contribute to the ability of MBONs to learn odor-specific responses. Lastly, we note that these learning rules are similar to homeostatic KC plasticity in that both provide a mechanism for increasing specificity of odor learning: the homeostatic model does so by eliminating broadly tuned KCs, the learning rule does so by allowing MBONs to ignore them.

While the odor-specific learning capacity of our mushroom body model is below the theoretical ideal, the KC representation of odors might still be “good enough” for associative learning in short-lived insects. It is also worth considering the extent to which odor-specificity is desirable: generalization of a learned association could be beneficial if the generalized odors have similar behavioral significance, or in the case of generalizing across multiple concentrations of an odor. The fact that honeybees show generalization of a conditioned response to odors that smell similar (to humans) to a trained odor, suggests that associative odor learning does generalize in insects^39-41^. In our model model, learning generalization (and our related measure of odor pair Intersection Fraction (IF)) was especially low between pairs of narrow odors. This suggests that odor receptor specificity could be a way for the nervous system to tune the extent of generalization by the mushroom body, with broadly tuned receptors producing more generalization, and narrowly tuned receptors producing odor-specific learned associations. Interestingly, in addition to the odorant receptor (OR)-expressing ORNs investigated in this study, insects express a second class of olfactory receptors called ionotropic receptors (IRs)^42^ that are narrowly tuned for specific amines and acids. While we do not have sufficient data to include IR-expressing ORNs in this model, the weak responses of OR-expressing ORNs to most amines and acids^1^ suggests that inclusion of IR-ORNs would lead the ORN representation of these odors to resemble that of other narrow odors in our tested ensemble. We would therefore expect lower rates of generalization between IR-activating amine or acid odor pairs, compared to odors that strongly activate the more broadly tuned OR population. This prediction is only made possible by modeling odor representations at the level of KCs, as odor-pair generalization is only weakly correlated with the correlation between odor representations at the level of PNs (**Fig S5**). Thus, in addition to compiling knowledge from a rich base of experimental data and providing a useful tool for investigations of odor representation and learning in insects, our model shows how the architecture of the mushroom body, the anatomy and dynamics of the olfactory periphery, and the tuning properties of olfactory receptors, all interact with each other to determine the ultimate form of associative odor learning and generalization.

## Methods

### ORN inputs to the dynamic PN model

#### Temporal dynamics of ORN responses

Model ORN responses were derived from a previously published experimental dataset^1^. Electrophysiological recordings in ORNs have shown that odor-evoked spiking undergoes little adaptation^25^, thus ORN firing rates in the dynamic model were assumed to be a simple step response convolved with the cell’s synaptic membrane filter (unless mentioned otherwise, all model cells were assumed to have a membrane time constant of 10 ms).

#### Generating synthetic ORN responses

ORN activation by odor mixtures was assumed to be a weighted sum of measured responses to single odors, with weights determined by each odor’s concentration in the mixture.

To study dimensionality of odor representations in different stages of olfactory processing, an additional 4890 “synthetic” odors were generated. For each synthetic odor, the response of each of the 23 simulated ORNs was randomly drawn from the set of that ORN’s responses to the 110 experimentally tested odors.

### Construction of the dynamic PN model

The dynamic PN model was built in two stages. First, we constructed a “static” model of the time-averaged PN response to a 500 ms odor presentation. This model is essentially the same as the model of Olsen et al, except for a change made to the form of the PN response linearity to allow PNs to drop below their spontaneous firing rate. From the static model, we obtain a set of four parameters determining the shape of the ORN-PN nonlinearity, and the strength of lateral inhibition between PNs. The dynamic PN model uses these same parameters, but introduces a dynamic model of GABA release and binding to GABA-A/B receptors to determine the time course of PN lateral inhibition. Spiking of the dynamic PN model is generated by an inhomogeneous Poisson process with an added refractory period, fit to match experimentally observed trial-to-trial variability in PN spike counts.

#### Static PN model: fitting the ORN-PN nonlinear transformation

The nonlinear input-output transformation from ORNs to their cognate PNs was modeled as in Olsen et al^2^, however to allow for odor-induced inhibition of PNs, we replaced the hyperbolic ratio function from the original paper with a thresholded hyperbolic tangent function: *PN*_evoked_ = [*PN*_spont_ + *R*_max_ tanh(*g* (*ORN*_evoked_ − *ORN*_spont_) + *c*)]_+_, where *X*_evoked_ and *X*_spont_ reflect the spontaneous and odor-evoked firing rates of PNs or ORNs, and *R*_max_, *g*, and *c* are fit parameters (see below). Spontaneous firing rates of ORNs were obtained from the Hallem and Carlson dataset. While PNs are also known to show spontaneous spiking at several Hz, characteristic spontaneous firing rates for each PN have not been published; these were instead generated from model parameters (see section on Spontaneous PN Activity below.)

To find values of parameters *R*_max_, *g*, and *c* that best reproduced the nonlinearity of the original model, we used nonlinear least squares to minimize the difference between the two models:

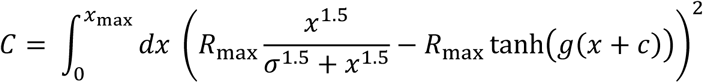

where *x* = *ORN*_evoked_ − *ORN*_spont_ and *x*_max_ is the largest experimentally-observed ORN response, and the value of *R*_max_ was kept the same in the original and revised model.

#### Static PN model: adding lateral inhibition

Lateral inhibition between glomeruli is mediated by a population of GABAergic local neurons (LNs) in the antenna lobe. Although LNs show heterogeneous morphologies and odor preferences, the LN population as a whole shows essentially the same spatial pattern of activation regardless of odor identity^24^.

Olsen et al found that the net effect of the LN population is to divisively normalize PN responses by the total ORN input to the antenna lobe. We kept this approach in our model, modeling LN-mediated inhibition of PNs with a divisive inhibition term:

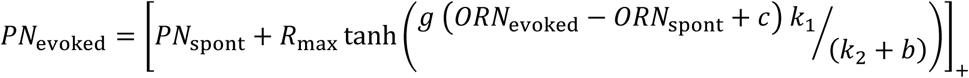

where 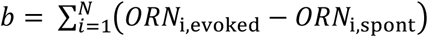 is the total ORN input to the antenna lobe, and *k*_1_ and *k*_2_ were again fit by nonlinear least squares to minimize the difference between models:

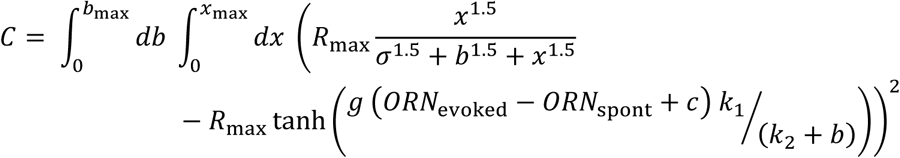

where *x* and *x*_max_ are defined as previously, and *b*_max_ is the maximum value of *b* computed from the Hallem and Carlson dataset.

#### Spontaneous PN activity

For the PN’s threshold (the value of (*ORN*_evoked_ − *ORN*_spont_) at which *PN*_evoked_ > 0) to be independent of *b* in our model, *PN*_spont_ should be proportional to *k*_1_/*k*_2_ – we therefore set *PN*_spont_ = *ORN*_spont_ *k*_1_/*k*_2_. In the fit model, this gave model PNs a mean spontaneous firing rate of 14.5 Hz, compared to 13.3 Hz measured in ORNs. Complete data on the spontaneous firing rates of PNs is unavailable, but they are known to show spontaneous activity of several Hz; the value used here can easily be adjusted in future models if additional information about PN responses becomes available.

#### Dynamic PN model: fitting slow time constants of inhibition

PN responses are shaped by slow lateral inhibition mediated by GABAergic LNs. As in previous models, we assumed the model LN population received equal input from all ORNs and projected onto all PNs with identical synaptic weights^2^, as well as feeding back onto themselves^21^. GABA concentrations at synaptic terminals was assumed to be a linear function of the LN firing rate, and GABA was taken to divisively inhibit both LNs and PNs, as described in previous studies^2,22^.

Previous experimental work^21^ found that GABA-induced IPSPs in LNs have a decay time constant *τ* of 100 ms and are completely abolished by picrotoxin, suggesting they are mediated by GABAA, while IPSPs in PNs are a sum of a fast-decaying (*τ* = 100 ms) picrotoxin-abolished component and a slow-decaying (*τ* = 400 ms) component that was abolished by GABAB antagonist CGP54626, suggesting a combination of GABAA- and GABAB-mediated inhibition in PNs. The dynamics of GABAA and GABAB receptor activation were modeled as

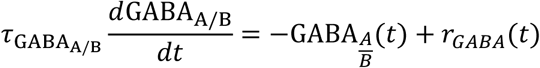

with 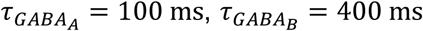, and where *c*_*GABA*_(*t*) is the extracellular concentration of GABA. As justified in the static model, we modeled the LN population as a single unit receiving excitatory input from all ORNs, and assumed extracellular GABA for all antenna lobe neurons to be a threshold-linear function of the LN firing rate, *c*_*GABA*_(*t*) = [*LN*(*t*)]_+_.

#### Dynamic model: GABA-mediated inhibition of LNs and PNs

The amount of GABAergic inhibition of PNs and LNs (I_PN_ and I_LN_, respectively) is a function of the relative levels of GABAA and GABAB receptor expression in the two populations. As LNs only appear to express the GABAA receptor (see previous section), we set *I*_*LN*_ = GABA_A_. In PNs, which express both receptor types, *I*_*PN*_ = *w*_*A*_GABA_A_ + *w*_*B*_GABA_B_; based on the analysis of GABA-induced IPSPs in Figure 2 of Wilson and Laurent^21^, we used *w*_*A*_ = 0.25 and *w*_*B*_ = 0.75.

We describe the firing rate of each PN in the dynamic model as a threshold-linear function of its inputs:

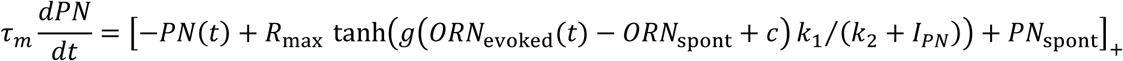

In fitting the model, we found that superlinear recruitment of LNs by the ORNs was needed for PN responses to match time-averaged firing rates predicted by the static model. Additional experimental investigation would help to further constrain the LN model and determine whether this feature is reasonable. The firing rate of the LN population in the dynamic model is given by:

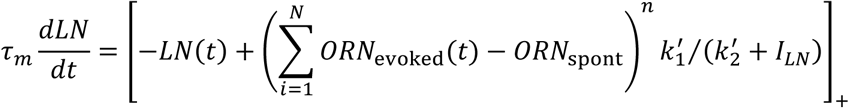

#### Parameter fitting in the dynamic model

Most parameters defining the PN nonlinearity and the effect of LN inhibition (*g, c, k*_1_, and k_2_) were kept the same as in the static model. One exception was the value of *R*_max_, which in the static model determined the maximum number of spikes fired during a one second stimulus presentation. Because most PNs transiently exceed their steady-state firing rate at odor onset, we increased *R*_max_ to 200Hz, the approximate peak firing rate observed in PNs.

To match the data to which the Olsen model was fit, the response of the dynamic model is computed as the average firing rate over a 500 ms odor presentation, minus the average spontaneous firing rate in a 500 ms window prior to odor presentation. The dynamic model is a good fit to the output of the Olsen model over all tested odors.

In the LN model, choice of *k*^′^_1_ and *k*^′^_2_ did not strongly affect PN performance, and for simplicity were chosen to be similar to *k*_1_ and *k*_1_. *m* and the cubic exponential were fit by hand to the Olsen model.

#### Spike generation from the dynamic PN model

To generate spikes from the dynamic model, instantaneous firing rates of model PNs were fed into a Poisson process with a 3 ms refractory period; the refractory period was fit to match the mean/variance relationship of PN spiking reported in ^2^. Five PNs were modeled per glomerulus, each with the same underlying firing rate but a different realization of the Poisson process.

### Construction of the spiking Kenyon cell model

Kenyon cells (KCs) receive excitatory input from a small number of PNs, and inhibitory input from the giant GABAergic anterior paired lateral (APL) interneuron. We modeled each KC as a leaky integrate-and-fire neuron, with membrane potential dynamics 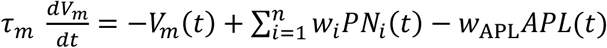 and spiking threshold θ. Three components of this model were derived from experimental data: the statistics of PN-KC connectivity, the dynamics and strength of APL inhibition onto KCs, and the spiking threshold.

#### PN-KC connectivity

PN-KC connectivity was derived from the experimental data of Caron et al^16^, who experimentally measured connection probability between PNs and KCs. Caron et al determined that PN-KC connectivity is random, in the sense that one KC claw forming a synapse with a given PN does not affect connection probabilities of the remaining claws. However, some PNs are much more likely to synapse with KCs than others.

For each model KC, the number of dendritic claws was randomly drawn from a set of 200 experimentally-measured claw counts (mean 6.8 claws, with max of 11 and min of 2.) PN inputs at each claw were randomly and independently drawn from a set of 342 measured connections between glomeruli and KCs; glomeruli in the set averaged 13.8 connections to KCs, with a max of 33. One glomerulus, da3, had no observed connections with KCs; other than that, the lowest number of observed connections was 2. After selecting a glomerulus, one of the five model PNs from that glomerulus was randomly selected as input to the given KC claw. PN-KC synaptic weights were assumed to be uniform, and normalized by the number of claws on each KC.

#### Kenyon cell spiking threshold: “full model”

Unlike PNs, KCs show little to no spontaneous activity: α, β, and β KCs are silent in the absence of odor, while α’ and β’ KCs have a spontaneous firing rate of around 0.1 Hz^3^. To match this in the model, we set each KC’s threshold to be a fixed amount above the level of depolarization produced by spontaneous input from model PNs, so that on average 10% of cells responded to any given odor from the Hallem and Carlson dataset.

The sparsity of odor-evoked responses in model KCs is determined both by the KC spiking threshold and by the strength of recurrent inhibition of KCs by the Anterior Paired Lateral (APL) neuron (see below). Lin et al found that the number of KCs responding to an odor roughly doubles when APL is silenced^30^. We thus set the KC threshold to achieve an average of 20% of KCs active across a panel of odors, then adjusted APL inhibition until the size of the responding KC population was halved to 10%.

#### Keyon cell spiking threshold: “homeostatic model”

In the homeostatic model, rather than setting a global threshold for the model KC population, we set the thresholds of individual KCs. For a given model KC, we calculated the PN input to that cell for a sample panel of 36 odors. For each odor, we computed the peak PN input to the model KC over a single trial (a 0.5-second odor presentation).

For the model without APL inhibition, we set the model KC’s threshold such that only the four odors with the highest peak inputs were above that threshold, thus achieving approximately 10% sparsity across odors. For the model with APL inhibition, we set the model KC threshold such that the seven odors (∼20% sparsity) with highest peak inputs were above that threshold, then adjusted APL inhibition until the size of the responding KC population was halved to 10%.

#### APL inhibition of Kenyon cells

APL is a giant interneuron that innervates the entire mushroom body, and releases GABA in a graded manner proportional to the number of KCs activated by an odor^29^. It extends a putative dendrite-like process to the lobes of the mushroom body, where KC axons are found, and an axon-like process to the calyx, where PN axons synapse with KC dendritic claws. It is thus believed that APL receives excitatory input from KC axons, and recurrently inhibits KCs either pre- or post-synaptically at their claws in the calyx.

In a previously published model of the mushroom body^20^, APL was assumed to decorrelate KC representations by removing the first principal component of PN responses in the 110-odor ensemble. While this approach substantially improved mushroom body performance on odor learning tasks, it resulted in patterns of KC activity which did not match well with available data. Specifically, the model predicted that public odors, like isopentyl acetate, would activate far fewer KCs than private odors like methyl salicylate—while the opposite is in fact the case.

We modeled APL release of GABA as a linear function of its membrane potential, after the results of Papadopoulou et al^29^, and assumed APL to inhibit KCs presynaptically, producing divisive inhibition. Because there is little data available regarding the strength of APL-KC synapses, our model APL receives equal input from all KCs, with APL membrane potential evolving as 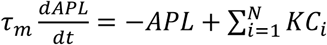. Similarly, we assumed APL formed equal-strength synapses onto all KCs in the population, with synaptic weight set such that APL activation approximately halved the number of active KCs, as described in the previous section.

### Measures of model performance

#### Inter-ORN/PN correlation

Defining *r*_*i*_(*n, t*) to be the response of the *i*^*th*^ PN to odor *n* at time *t*, and 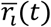 to be the average response of that PN across all odors at time *t*, **the inter-PN correlation** between PNs *i* and *j* is defined as 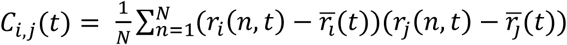. Conversely, given two odors *n* and *m*, the **inter-odor correlation** is defined as 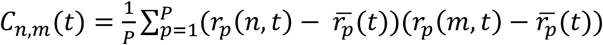.

#### Lifetime sparseness

Lifetime sparseness is defined as in^13^, and is derived originally from^43,44^. For individual KCs or PNs (as in **Fig 4A-B**), lifetime sparseness is given by 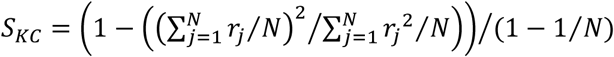, where *N* is the number of odors and *r*_*j*_ is the response of the KC to odor *j*. Lifetime sparseness of an odor across the KC population is defined similarly, replacing *N* with the number of neurons (rather than odors) and *r*_*j*_ with the response of neuron *j* to the odor in question.

#### KC intersection fraction

Given set A of KCs that responded to odor *a*, and set B of KCs that responded to odor *b*, the intersection fraction” (IF) is defined for odor *a* as |*A* ∩ *B*|/|*A*| (or for odor *b* as |*A* ∩ *B*|/|*B*|). This measure therefore gives the fraction of KCs responding to a given odor that also respond to a second odor. Or, if we consider a learning algorithm that potentiates KC-MBON synapses of all KCs responding to odor *b*, this measure tells us what percentage of odor-*a*-responding KCs will have potentiated synapses after learning.

We selected this measure over other dissimilarity metrics because the IF of odor *a* with odor b is a reasonably accurate predictor of the probability that a single-class Perceptron algorithm trained to respond to odor *b* will also respond to odor *a* (**Fig S5**). We computed this value for all pairs of odors in our 110-odor dataset to estimate the distribution of odor overlaps in different version of the mushroom body model.

#### Broad vs narrow odors

We define broad odors as those that strongly activate many ORNs, and narrow odors as those that strongly activate only one or a small number of ORNs. To identify the “narrowest” odors in our 110-odor dataset, we selected the odors with the greatest ratio between the maximum ORN response to that odor and the mean ORN response magnitude to that odor. To identify the “broadest” odors in our dataset, we selected those with the highest mean ORN response.

#### Perceptron learning and overgeneralization

This model is based on a single idealized neuron (here an MBON) that receives input from a population of *N* input neurons (our model KCs) via a set of synapses with weights *w*, and produces response *r*_*ON*_(*q*) = *θ*(*w* * *r*_*kc*_ (*q*)) where *r*_*kc*_ (*q*) is the vector of KC responses to odor *q, r*_*ON*_ is the firing rate of the MBON, and *θ* is the sign function.

Synaptic weights are allowed to fall below 0 in this model, however this could be prevented by adding a non-negativity constraint to *w* and setting *θ*(*x*) = sign(*x* − *ϕ*) for some threshold *ϕ*.

For single-class learning with a set of *Q* reinforcer-paired odors, we set the target value of *r*_*ON*_(*q*) to be −1 for odor *q* in *Q*. The target value of *r*_*ON*_(*q*) for all odors not in *Q* is 1, however these odors are not seen during training. We initialized all KC-MBON synaptic weights to 1, and trained weights using the standard perceptron algorithm with a learning rate of η=0.01 until *θ*(*w* * *r*_*kc*_(*q*)) = −1 for all *q* in *Q*, or for a maximum of 50,000 training iterations. Overgeneralization of this model is given by the fraction of unpaired odors *p* ∉ *Q* for which *θ*(*w* * *r*_*kc*_(*p*)) = −1. All measurements were averaged over 15 instantiations of the mushroom body model, and for each instantiation we tested 50 repeated samples of paired odor set *Q* from the full set of 110 simulated odors.

#### Synaptic learning and overgeneralization

Learning with the two-part anti-Hebbian rule 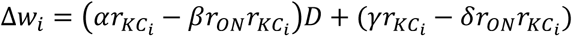 was simulated using parameters *α* = 20, *β* = 1, *γ* = 1, *δ* = 1. KC-MBON synaptic weights were initialized to random positive values, and learning was allowed to proceed until the mean of Δ*w* fell below a threshold or a maximum number of iterations was reached. Overgeneralization in the model was defined as the fraction of unpaired odors for which the MBON response was below the maximum MBON response to paired odors.

#### Other predictions regarding the dynamics of dopamine signaling during odor learning

In the text, dopamine provided a simple binary reinforcement signal, with D=1 whenever a reinforcement signal occurs, and D=0 otherwise. But it is possible that the amount of dopamine released in the mushroom body could also shape the dynamics of learning. Here we present two extensions of our model to explore possible implications of graded dopamine release.

First, two types of dopamine receptors are known to be expressed in the mushroom body: DAMB has a high affinity for dopamine, and hence a low threshold for activation^45^, while dDA1 has a low affinity for dopamine, and hence a much higher threshold for activation^46^. A simple modification of our two-part learning rule would be to assign each part to a separate receptor, as: 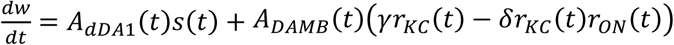, where *A*_*dDA*1_(*t*) and *A*_*DAMB*_(*t*) reflect the degree of activation of dDA1 or DAMB. This model behaves similarly to the form presented in the text, however it predicts that a complete silencing of dopamine neurons (or knockout of both dopamine receptors) would block both odor learning and odor habituation. Importantly, this model also suggests that it should be possible to separately block either the associative or the non-associative components of learning, by specifically knocking out either dDA1 or DAMB receptor. While knocking out dDA1 would impair associative learning of odor-reinforcer pairings (as has been shown^46^), a fly with a DAMB knockout would not show impaired associative learning, but would be expected not to habituate to odors, and to overgeneralize more quickly to unpaired odors.

Because dopaminergic neurons receive extensive feedback input from MBONs^8^, we also considered learning rules in which dopamine input takes the form of a prediction error, *D* = (*R*^*^ − *r*_*ON*_) for odors paired with reinforcement, and *D* = (*R*^0^ − *r*_*ON*_) for unpaired odors, where *R*^0^ and *R*^*^ are constants. In the simplest case where 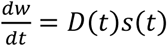, this rule has fixed points *r* = *R*^*^ for paired odors and *r*_ON_ = *R*^0^ for unpaired odors, although depending on parameter values one or the other fixed point may be replaced by *r*_*ON*_ = *α*/*β*. This rule predicts that dopaminergic neuron activity will change over the course of learning or habituation, eventually declining to zero or near-zero. As a result, stopping dopamine reinforcement will have no effect on MBON responses to paired or unpaired odors, in contrast to the prediction of our first model. This rule also predicts that MBONs will not show habituation to novel odors when dopaminergic neurons are silenced.

### Code Availability

Code for building and simulating all versions of the model and code for learning/generalization investigations is provided with documentation at github.com/annkennedy/mushroomBody.

## Acknowledgements

The author is grateful to L.F. Abbott, Richard Axel, Daisuke Hattori, Peter Wang, and Glenn C. Turner for many helpful conversations during the development of this model, and Elizabeth J. Hong and Vanessa Ruta for their comments and feedback during the preparation of this manuscript. The author was supported by postdoctoral fellowships from the Swartz Foundation and Helen Hay Whitney Foundation.

## Works cited

**Figure S1.**
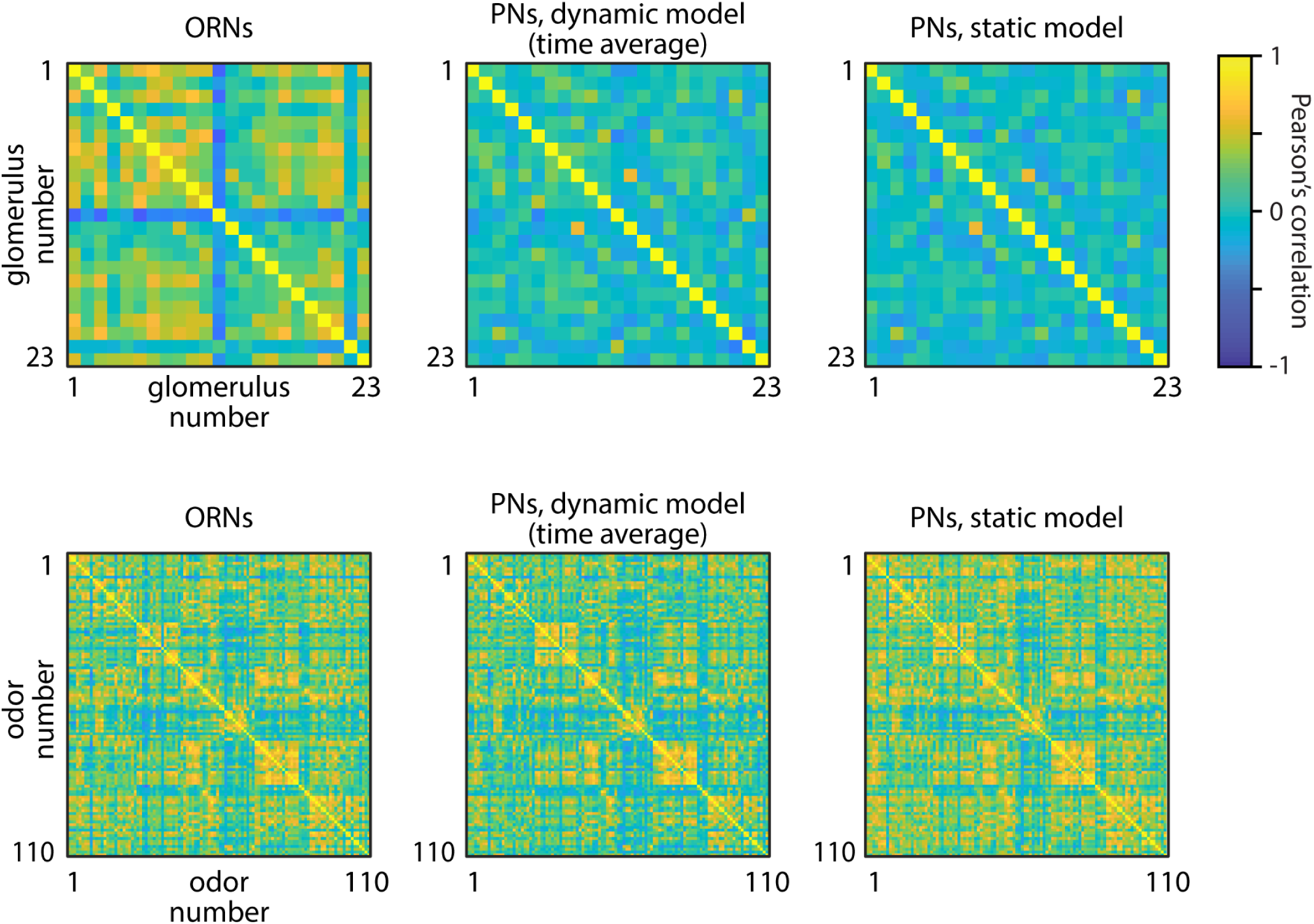
between-glomerulus vs between-odor correlations. Correlation matrices between glomeruli and between odors, in the ORN and PN populations. Top row, between-glomerulus correlation matrices, showing a drop in inter-glomerulus correlations from ORNs to PNs in both the time-averaged dynamic model and the static model, as reported previously^2^. Bottom row, despite a drop in inter-glomerulus correlations, the inter-odor correlations remain largely unchanged from ORNs to PNs.

**Figure S2.**
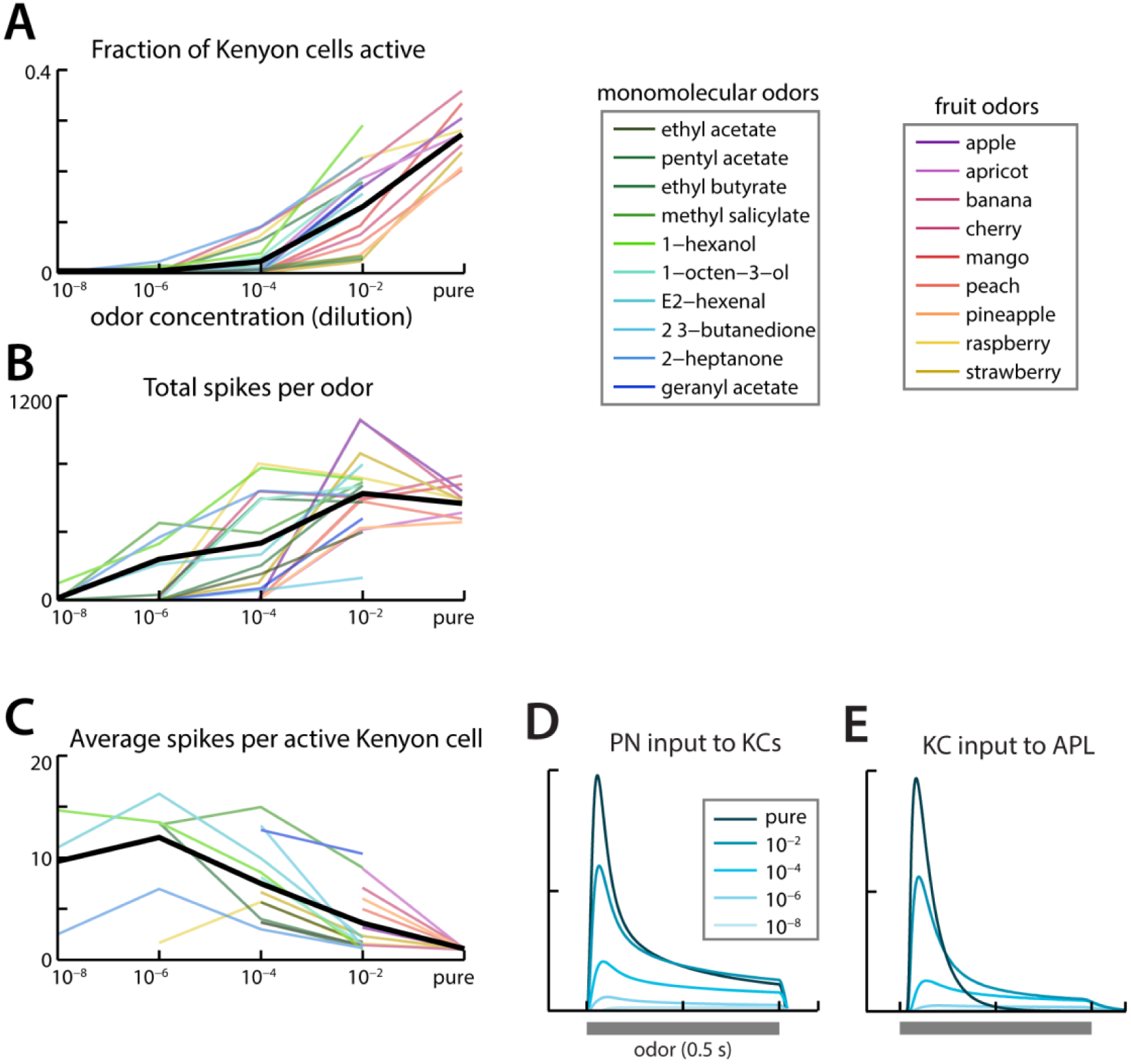
Odor-concentration dependence of model KC population. KC responses from the full model using ORN responses to multiple dilutions of monomolecular and fruit-based odorants, from Hallem and Carlson. **A)** The fraction of KCs responding to an odor increases with odor concentration, consistent with our expectations from experimental observations. **B)** The total number of KC spikes fired for odor shows a more modest dependence on odor concentration than the number of active KCs. **C)** The results in B and A are explained by the fact that as odor concentration increases, active model KCs reduce the number of spikes fired. **D-E)** Average dynamics of PN input to KCs (D) and KC input to APL (E), showing that an increase in odor concentration causes a higher peak in odor-evoked inputs to KCs and to APL. The stronger PN input to KCs leads to a greater number of KCs spiking at high odor concentrations, as seen in A. The increased recruitment of APL inhibition back onto KCs then inhibits active KCs, causing the KC representation of odors to become more transient and hence reducing the average number of spikes per KC at higher concentrations.

**Figure S3.**
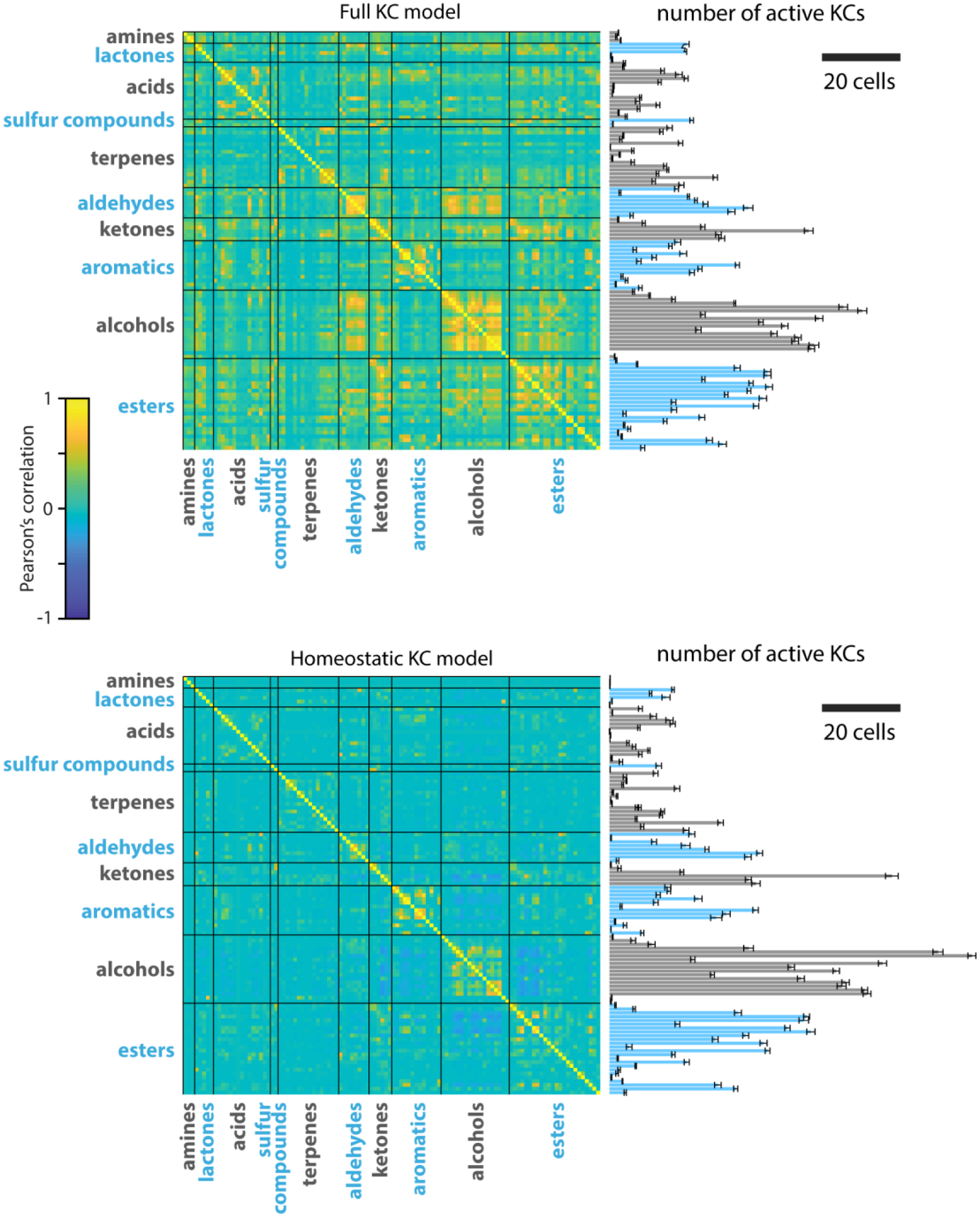
Correlation matrices with odor classes. Correlation matrices between odors in the model KC population, for both the full model (top) and the homeostatic model (bottom), with odor classes indicated. Odor classes are as in ^1^, see that paper for identities of all tested odors. Bar graphs show the number of model KCs responding to each odor (mean ± STD across 10 instantiations of each model). Note that different instantiations of the models all have very similar active KC counts for each odor, showing the KC responses can be quite reliable across individuals despite the random nature of PN-KC connectivity^47^.

**Figure S4.**
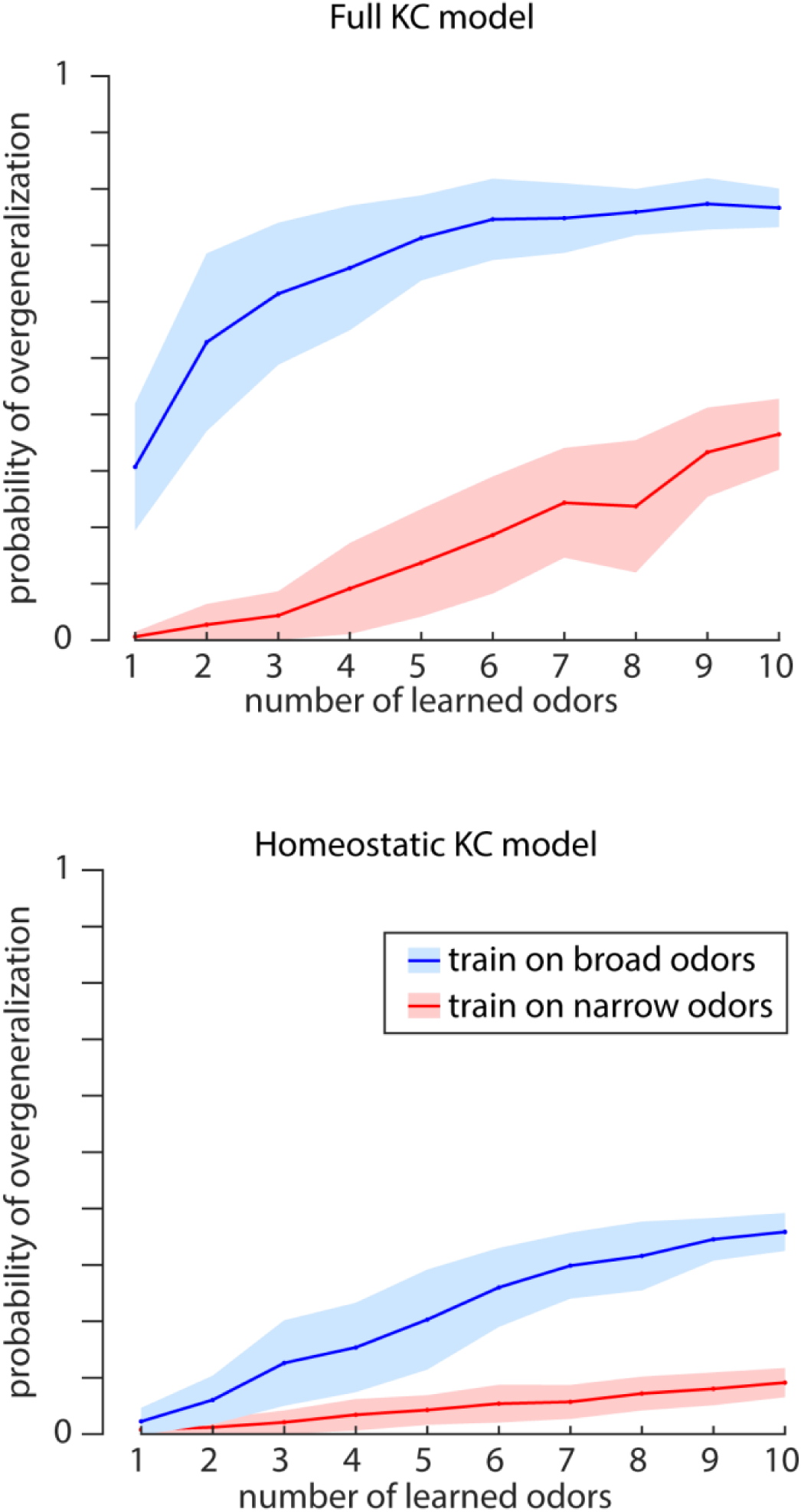
Single-class perceptron overgeneralization to broad vs narrow odors. Probability of single-class perceptron generalization when training odors were selected from only the 10 “broadest” odors or only the 10 “narrowest” odors, in the full KC model and the homeostatic KC model. See **Methods** for metric used to identify broadest vs narrowest odors.

**Figure S5.**
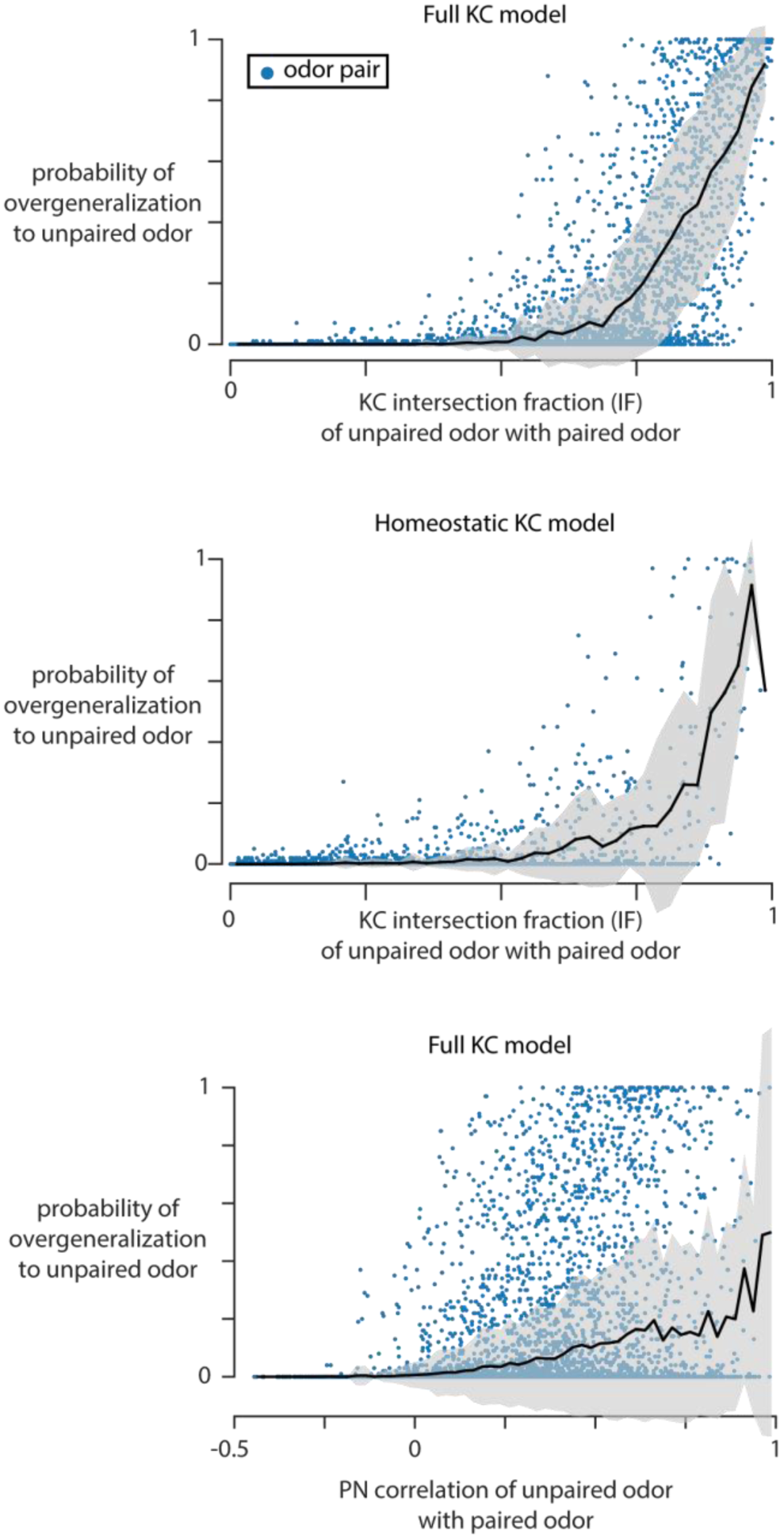
Single-class perceptron overgeneralization is well predicted by the intersection fraction (IF) with the paired odor. We computed probability that a single-class perceptron trained to recognize odor *a* (the “paired odor”) also produces a positive response to odor *b*, for all (*a,b*) odor pairs in the 110-odor dataset. **Top+middle**, this value is plotted against the intersection fraction (IF) of odor *b* with odor *a*, defined as the fraction of odor *b* responding neurons that also respond to odor *a*; **bottom** this value is plotted against the Pearson’s correlation between odors *a* and *b*. We find that the IF is a better predictor of perceptron overgeneralization than the correlation between odor representations at the PN or KC level (KC correlations not shown). Black lines and envelopes show mean +/− STD probability of overgeneralization across odor pairs in a 0.025-wide window along the x-axis.

